# The microtubule-associated protein CLASP is translationally-regulated in light-dependent root apical meristem growth

**DOI:** 10.1101/2020.04.20.051417

**Authors:** Laryssa Halat, Katherine Gyte, Geoffrey Wasteneys

**Affiliations:** Department of Botany, The University of British Columbia, 6270 University Blvd, Vancouver, BC, V6T 1Z4, Canada

**Keywords:** Meristem, CLASP, microtubule, hormones, etiolation

## Abstract

The ability for plant growth to be optimized, either in the light or dark, depends on the intricate balance between cell division and differentiation in specialized regions called meristems. When *Arabidopsis thaliana* seedlings are grown in the dark, hypocotyl elongation is promoted, whereas root growth is greatly reduced as a result of changes in hormone transport and a reduction in meristematic cell proliferation. Previous work showed that the microtubule-associated protein CLASP sustains root apical meristem (RAM) size by influencing microtubule (MT) organization and by modulating the brassinosteroid (BR) signalling pathway. Here, we investigated whether CLASP is involved in light-dependent root growth promotion, since dark-grown seedlings have reduced RAM activity that is observed in the *clasp-1* null mutant. We showed that CLASP protein levels were greatly reduced in the root tips of dark-grown seedlings, which could be reversed by exposing plants to light. We confirmed that removing seedlings from the light led to a discernible shift in MT organization from bundled arrays, which are prominent in dividing cells, to transverse orientations typically observed in cells that have exited the meristem. BR receptors and auxin transporters, both of which are sustained by CLASP, were largely degraded in the dark. Interestingly, we found that despite the lack of protein, *CLASP* transcript levels were higher in dark-grown root tips. Together, these findings uncover a mechanism that sustains meristem homeostasis through CLASP, and advances our understanding of how roots modulate their growth according to the amount of light and nutrients perceived by the plant.

**One Sentence Summary:** The microtubule-associated protein CLASP is regulated at the translational level when root meristem growth is inhibited in dark-grown plants.

## INTRODUCTION

At the beginning of their lives, plants face a precarious situation. Resources stored in the seed will only sustain life for a limited time, and thus it is critical that plants quickly emerge from the soil to begin the process of photosynthesis and sugar production. Plants must therefore use their initial resources strategically during the first days of development to ensure survival. It is known that when seedlings are germinated and grown in complete darkness (skotomorphogenesis), the hypocotyl grows rapidly, whereas root elongation is inhibited. When grown in light conditions (photomorphogenesis), sucrose derived from aerial organs is sufficient to promote root elongation in *Arabidopsis* (Kircher and Schopfer, 2012), and to reduce hypocotyl growth. One fundamental question is how growth is promoted in some organs and inhibited in others to meet the needs of a plant during developmental transitions.

Several signalling pathways are necessary for plants to withstand stress due to lack of nutrients, which is the case in extended periods of darkness. Autophagy is a conserved degradation pathway in eukaryotes that is activated during cellular starvation. When glucose is available, the formation of autophagosomes is inhibited by increased reactive oxygen species (Huang et al., 2018). Several proteins are targeted for degradation through the target of rapamycin (TOR) kinase, which is a highly conserved eukaryotic protein that integrates environmental signals with downstream developmental and metabolic pathways, such as translation, protein degradation, and cell division (reviewed in Dobrenel et al., 2016). Previous work has found that sugar, which activates TOR, inhibits autophagy-mediated degradation of the BZR1 transcription factor that normally regulates thousands of genes through the brassinosteroid (BR) signalling pathway. In times of cellular starvation and growth arrest, TOR is inactive and BZR1 is consequently degraded (Zhang et al., 2016). Similarly, a related BR transcription factor BES1 forms a complex with DSK2 and AT8G, which directs cargo to autophagosomes to be degraded via autophagy in times of stress (Nolan et al., 2017). Thus, it is clear that plants respond to stress through crosstalk of BR and autophagy signalling pathways, but these mechanisms likely differ between hypocotyls and roots to control organ-specific growth.

Research into the function of Cytoplasmic Linker Associated Protein (CLASP) revealed its pivotal function in maintaining root meristem size through control of microtubule (MT) organization. Mutants that lack CLASP are dwarf, with fewer cells in the root division zone, indicating a role for CLASP in cell division (Ambrose et al., 2007). In plants, CLASP is involved in anchoring MTs to the cell cortex and is considered to be a stabilizing factor for MTs (Ambrose and Wasteneys, 2008). Through confocal imaging, CLASP has been found to distribute to the sharp transverse edges of newly-divided cells in the RAM, which normally present a barrier to growing MTs and cause them to undergo catastrophe upon encounter (Ambrose et al., 2011). The presence of CLASP at these edges, however, enables formation of transfacial MT bundles (TFBs) that span the periclinal and transverse faces of the root meristematic cells. TFBs are associated with maintaining the capacity for cell division and are not present in elongating cells. Recent work has concluded that without CLASP, and consequently TFBs, meristem size is reduced because cells prematurely start to elongate instead of continuing to divide.

Two recent studies demonstrated CLASP’s involvement in both auxin and BR hormone signalling. CLASP directly interacts with the retromer component sorting nexin 1 (SNX1) and tethers SNX1-associated endosomes to MTs (Ambrose et al., 2013). Stabilization of SNX1 along MTs fosters recycling of the auxin efflux carrier PIN2 to the plasma membrane (PM), thereby reducing PIN2 transport to vacuoles for degradation (Ambrose et al., 2013). In mutants lacking *CLASP* expression, PIN2 is depleted at the PM, resulting in auxin accumulation in the root tip, consistent with PIN2’s function in directing auxin away from the quiescent centre via the epidermis and cortex tissues (Ambrose et al., 2013). The CLASP-SNX1 interaction also promotes recycling of the BR receptor BRI1 to the PM, thus enhancing BR signalling. Ruan et al. (2018) identified a negative feedback loop whereby the BR-activated transcription factors BZR1 and BZR2/BES1 bind to the *CLASP* promoter and repress *CLASP* gene expression. The downregulation of *CLASP*, through application of exogenous BR or in mutants with constitutively active BR signalling, is strongly correlated with premature exit of cells from the division zone, producing a smaller meristem phenotype similar to that observed in *clasp-1* mutants.

To determine how root growth is regulated in early plant development, we compared the role of CLASP in actively proliferating light-grown meristems to those grown in dark conditions when meristem growth is largely inhibited. Notably, in the absence of BZR1 in the dark, *CLASP* transcript levels were elevated yet despite this, CLASP protein levels were greatly reduced. In addition, many CLASP-dependent processes such as MT organization and hormone signalling pathways were disrupted. Our work reveals that CLASP is regulated post-transcriptionally based on light signals that control when root growth is desirable for a plant.

## RESULTS

### CLASP is required for increased cell proliferation in response to light and/or sucrose

To investigate CLASP’s involvement in light-dependent root meristem activity, we compared *clasp-1* null-transcript mutants and wild-type root growth in light and dark conditions. Although it is well known that hypocotyl expansion is stimulated and that root growth is inhibited in the dark, we noted that most previous dark-growth investigations with the model system *Arabidopsis thaliana* included some sucrose in the media. Considering the likelihood that sucrose, a product of photosynthesis, signals successful germination and stimulates root growth, we compared organ growth in culture media either lacking or supplemented with sucrose.

Root growth responses indicated that CLASP’s function is strongly associated with the rapid root growth that is stimulated in the light. Whereas light stimulated a 7-fold length increase in 6 day-old wild-type roots from 5.23 ± 0.40 mm to 37.50 ± 3.03 mm, it only caused *clasp-1* root length to double from 3.47 ± 0.24 mm to 7.33 ± 0.94 mm (Fig. 1A, C). We also noted that dark-grown wild-type roots were of similar length to *clasp-1* mutants, suggesting that dark-grown wild-type meristems are deficient in CLASP.

**Figure 1.**
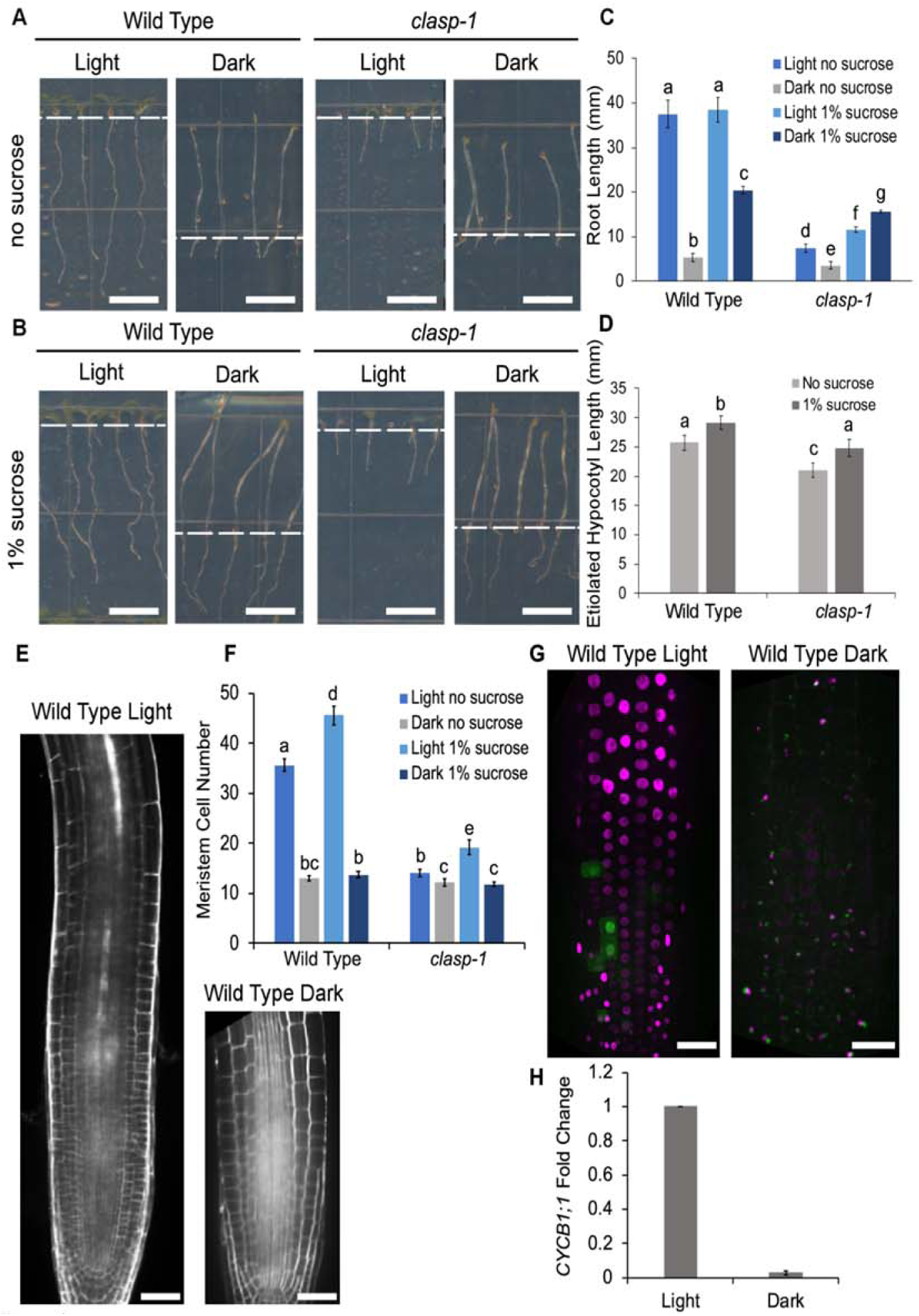
Light and sucrose control growth of the root apical meristem A, Seedling morphology of wild type and *clasp-1* mutants grown in either 24h light or 24h dark conditions for 6 days with no sucrose. Scale bar = 0.5 cm. B, Seedling morphology of wild type and *clasp-1* mutants in light and dark growth conditions, grown in the presence of 1% sucrose. Scale bar = 0.5 cm. The dashed white lines in A and B represent the boundary between roots and hypocotyls. C, Quantification of root lengths in the seedlings depicted in A and B. Different letters indicate significance (p<0.01) assessed by a two-way ANOVA with interaction. n = 30 for each growth condition. D, Quantification of etiolated hypocotyl lengths in dark-grown seedlings depicted in A and B. Different letters indicate significance (p<0.01) assessed by a two-way ANOVA with interaction. n = 30 for each growth condition. E, Wild type root meristems stained with propidium iodide to show cell outlines. Scale bar = 30 μm. F, Quantification of meristem cell number in wild type and *clasp-1* roots without sucrose in light and dark growth conditions. Different letters indicate significance with a two-way ANOVA. n = 30 for each genotype and growth condition. G, Confocal z-projection of Cell Cycle Tracking in Plants (CYTRAP)-labelled roots grown in the light or dark. Nuclei lacking fluorescence are in G1, magenta-labelled nuclei are cells in S phase, and green fluorescent nuclei represent cells undergoing mitosis. Scale bar = 30 μm. H, qRT-PCR of *CYCB1;1* expression fold change in light and dark-grown root tips. *MUSE3* was used as a reference gene. Error bars indicate SEM. n = 3 biological replicates.

We next determined that supplementing the culture medium with a moderate (1% weight/volume) amount of sucrose caused a major increase in the length of dark-grown roots, but that this effect was independent of CLASP. Treatments with 1% sucrose increased dark-grown wild-type root lengths from 5.23 ± 0.40 mm to 20.42 ± 0.88 mm, and increased mean *clasp-1* root lengths from 3.47 ± 0.24 mm to 15.59 ± 0.83 mm (Fig. 1B, C), which is an approximate 4-fold increase for both genotypes. This indicates that the sucrose-dependent growth stimulation is CLASP-independent. By comparison, sucrose had no significant effect on root length for light-grown wild-type seedlings and a moderate increase (from 7.33 ± 0.94 mm to 11.56 ± 0.60 mm) for *clasp-1*.

Based on the fact that hypocotyl expansion is entirely dependent on cell elongation, we compared the etiolated hypocotyl responses of dark-grown wild-type and *clasp-1* mutants to sucrose. Although *clasp-1* hypocotyls were significantly shorter than those of wild type under both conditions, *clasp-1* and wild-type hypocotyls responded in the same manner to 1% sucrose, with an approximate 10% length increase (Fig. 1D). These findings demonstrate that although sucrose has relatively little effect on dark-grown hypocotyls, the increase is CLASP independent.

The observation that dark-grown wild-type and *clasp-1* mutants have similar root lengths prompted us to investigate whether the light-induced changes in root elongation can be attributed to CLASP-dependent changes in cell proliferation. Growing seedlings in the light more than doubled the number of cells in wild-type root meristems but only increased the number of cells in *clasp-1* mutant root meristems by 10% (Fig. 1E, F). Furthermore, the number of cells in dark-grown wild-type seedlings was similar to that found in *clasp-1* mutants (Fig. 1E, F). The addition of 1% sucrose to dark-grown seedlings did not affect cell proliferation in either genotype (Fig. 1F), suggesting that the increase in root length (Fig. 1C) was mainly due to cell elongation. These data indicate that the light-induced increase in root cell proliferation is largely dependent on CLASP.

To investigate the basis for light-dependent increases in cell proliferation, we used the Cell Cycle Tracking in Plants (Cytrap) fluorescent marker line to identify cells in S-phase (CDT1a-RFP, magenta) and mitosis (CYCB1-GFP, green) (Yin et al., 2014). Both reporters showed a dramatic loss of fluorescence in dark-grown roots without sucrose (Fig. 1G). Consistent with the lack of CYCB1-GFP protein (Fig. 1G), *CYCB1;1*, which is upregulated specifically in G2/mitosis, showed very low levels of expression in dark-grown root tips, as measured by quantitative-Real Time-PCR (qRT-PCR) (Fig. 1H).

The data shown in Figure 1 clearly demonstrate that the inhibition of root growth in the dark is associated with reduced cell proliferation, and that *clasp-1* knockout mutants are less affected by these conditions. Taken together, these findings indicate that CLASP mediates the cell proliferation that is stimulated when seedlings are exposed to light.

### The BR pathway is dampened in dark-grown roots, leading to an elevation of *CLASP* transcript levels

Previous work (Zhang et al., 2016) showed that dark-induced cellular starvation led to BZR1 degradation by autophagy. Using a BZR1-CFP fluorescent reporter, we confirmed that the BZR1 transcription factor accumulates in the nuclei of light-grown root cells with and without sucrose, but was absent when plants were grown in media lacking sucrose in the dark (Fig. 2A). By contrast, dark-grown roots with 1% sucrose in the media still showed BZR1-CFP in the nuclei (Fig. 2B). Since germinating seedlings grown in dark conditions do not yet have a source of sucrose, and the BR-activated transcription factor BZR1 was only degraded under these conditions, our remaining analyses focused on roots grown in the absence of sucrose.

**Figure 2.**
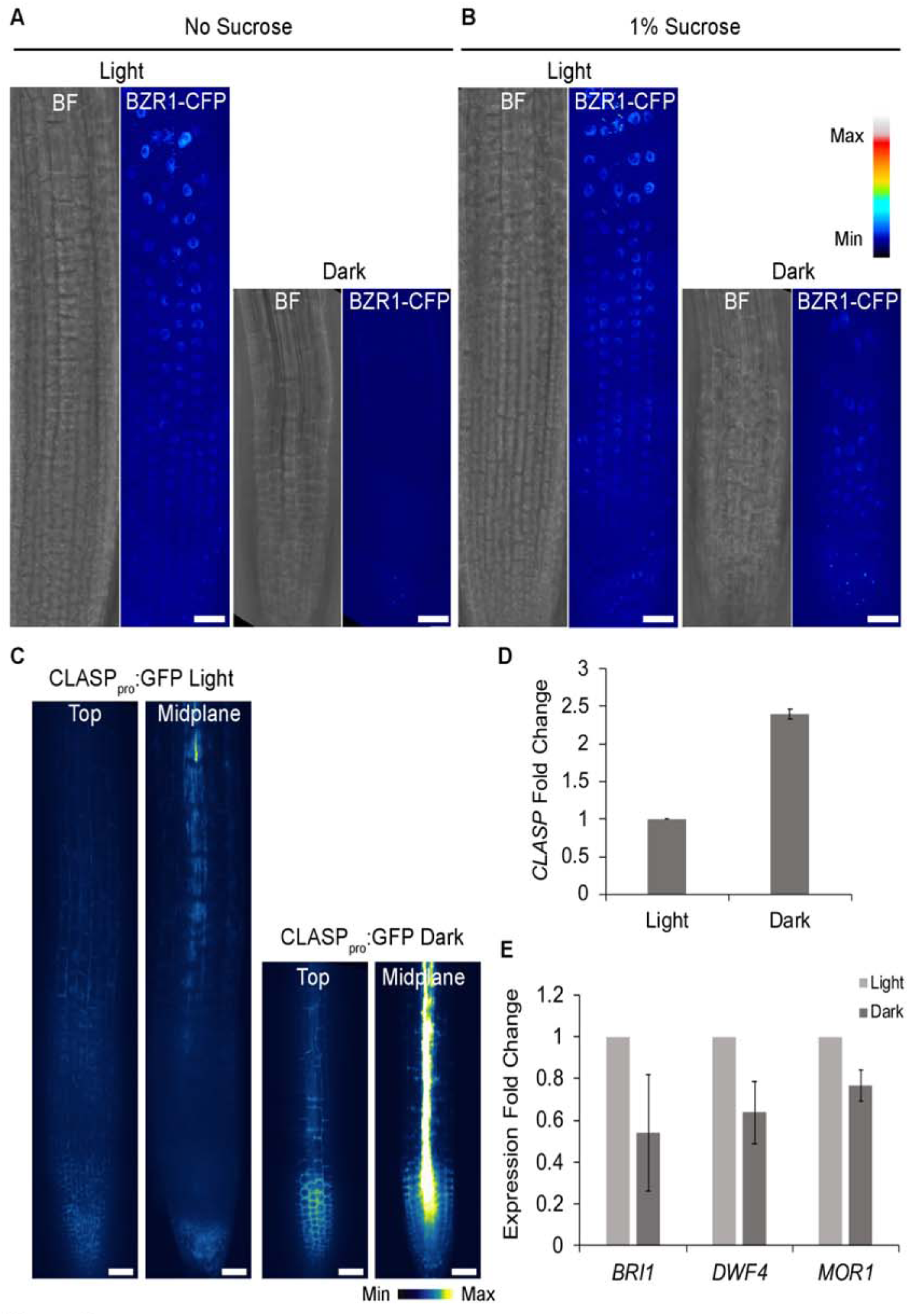
Brassinosteroid signalling is dampened in dark-grown root meristems A, B, Confocal z-projections of light and dark-grown root meristems grown with no sucrose (A) and 1% sucrose (B) showing brightfield (BF) and BZR1-CFP channels pseudo coloured with a lookup table. Scale bar = 30 μm. C, *CLASP*_*pro*_:*GFP* in light and dark-grown root tips. Confocal images are shown from the top (epidermal) layer of tissue as well as in the midplane of the root. Scale bar = 30 μm D, qRT-PCR analysis of *CLASP* expression in light and dark-grown root tips. *MUSE3* was used as a reference gene. Error bars indicate SEM. n = 3 biological replicates. E, qRT-PCR of two genes involved in the brassinosteroid pathway, *BRI1* and *DWF4*, and a microtubule-associated protein, *MOR1*, in light and dark-grown root tips. *MUSE3* was used as a reference gene. Error bars indicate SEM. n = 3 biological replicates.

Given that BZR1 is a negative regulator of *CLASP* transcription, we hypothesized that *CLASP* expression would be greater in dark-grown root meristems. Indeed, the *CLASP* promoter driving the expression of free GFP (*CLASP*_*pro*_:*GFP*) revealed greater fluorescence in dark-grown roots than in plants grown in the light (Fig. 2C). Interestingly, the level of free GFP in dark-grown roots was especially high in the stele (Fig. 2C). Using qRT-PCR from root-tip RNA, we found an almost 2.5-fold increase in *CLASP* gene expression in the dark (Fig. 2D), which is consistent with the observed increase in *CLASP*_*pro*_:*GFP* fluorescence.

Since BZR1 is a key effector of BR signalling, we predicted that other genes in the BR pathway would be differentially regulated in the dark, minus-sucrose conditions. Genes encoding the BR receptor *BRI1* and the BR-biosynthetic enzyme *DWF4*, had reduced expression levels by about 45% and 35% respectively (Fig. 2E), indicating a general loss of the BR signalling pathway in dark-grown roots. To determine if genes encoding other microtubule-associated proteins were affected in a similar manner to *CLASP*, we evaluated transcript levels of *MOR1*, which encodes a protein that binds to microtubule plus ends and has domain similarity to CLASP. qRT-PCR analysis showed that plants grown in the dark had a 25% decrease in *MOR1* expression (Fig. 2E). Thus, *CLASP* transcript levels are higher in the absence of BR signalling when roots are exposed to dark conditions, and this pattern is not observed for *MOR1*, indicating that the increased *CLASP* expression is a direct consequence of the loss of BR signalling.

### CLASP protein levels are diminished under dark conditions

The high level of *CLASP* transcript in dark-grown roots is at odds with the shutdown of meristem activity. Since it is impossible to extract sufficient protein from the root apical meristem for immunoblotting, to assess CLASP protein levels we used a GFP-CLASP fluorescent reporter driven by its endogenous promoter (Ambrose et al., 2011). In light-grown root meristems, GFP-CLASP was abundant along the transverse edges of cells in the division zone (Fig. 3A). In dark-grown root meristems, by contrast, only a few GFP-CLASP puncta were observed around the cell edges (Fig. 3B), and fluorescence intensity was reduced by about half (Fig. 3C). It is important to note that autofluorescent bodies were also observed in dark-grown root cells using microscope settings equivalent to those used for GFP fluorescence (Supplemental Fig. S1) and should not be confused with GFP-CLASP puncta seen in Fig. 3B. To further investigate the light-dependence of GFP-CLASP protein levels, we transferred plants grown in dark conditions to growth chambers with 24h light, and vice versa (Fig. 3D, E). In plants transferred from dark to light (Fig. 3D), GFP-CLASP protein levels increased over a period of 4 days, and gradually disappeared over the same time period in plants transferred from light to dark (Fig. 3E). These results indicate a clear disconnection between light-dependent control of *CLASP* transcription and CLASP protein levels.

**Figure 3.**
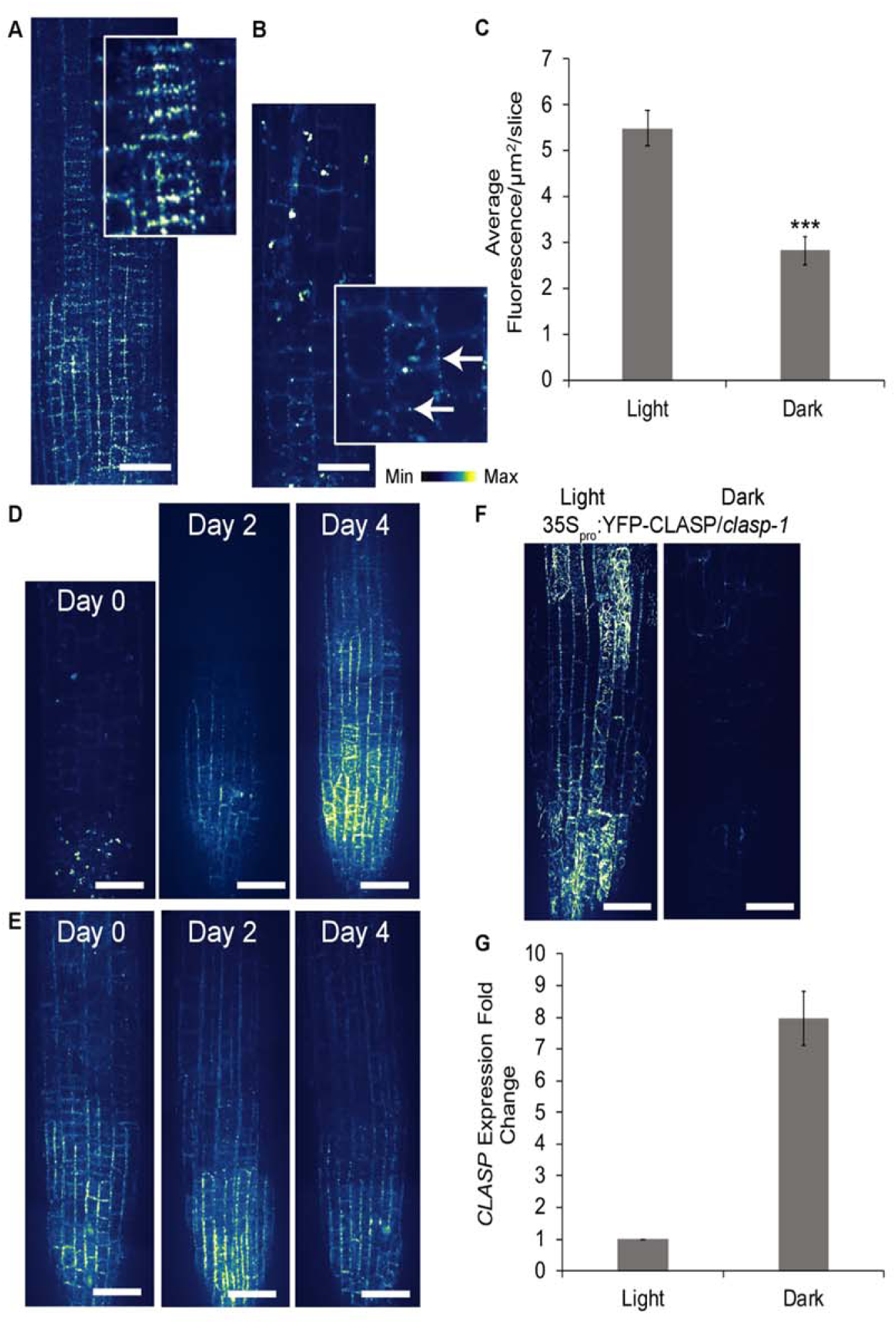
CLASP protein levels are diminished in dark-grown root meristems A and B, Confocal z-projection of root division zone epidermal cells expressing *CLASP*_*pro*_:*GFP-CLASP* grown in the light (A) and dark (B) for 6 days. The insets show magnified views of GFP-CLASP on cell edges. Scale bar = 30 μm. C, Quantification of fluorescence intensity in the root division zone of *CLASP*_*pro*_:*GFP-CLASP/clasp-1* expressing plants. *** p<0.001, Student’s t-test with equal variance. n = 27 roots for light and 26 roots for dark. D, Confocal z-projection of *CLASP*_*pro*_:*GFP-CLASP/clasp-1* plants germinated and grown in the dark for 4 days (Day 0) then transferred to 24h light conditions. Images show *CLASP*_*pro*_:*GFP-CLASP* 2 and 4 days following transfer to light. E, *CLASP*_*pro*_:*GFP-CLASP/clasp-1* plants germinated and grown in the light for 4 days (Day 0) then transferred to 24h dark conditions. Images show *CLASP*_*pro*_:*GFP-CLASP* 2 and 4 days following transfer to the dark. F, Confocal z-projection of *35S*_*pro*_:*YFP-CLASP* plants grown in the light and dark for 6 days. Scale bar = 30 μm. E, qRT-PCR of *CLASP* expression in *35S*_*pro*_:*YFP-CLASP* root tips grown in the light or dark for 6 days. *MUSE3* was used as a reference gene. Error bars denote SE; n = 3 biological replicates.

To further test whether translational control of CLASP is disconnected from transcriptional regulation, we used lines with *YFP-CLASP* driven by the 35S promoter to overexpress *CLASP* and compared root-tip transcript and protein levels in the light and dark. As shown in Fig. 3F, although *35Spro:YFP-CLASP* had greatly increased fluorescence in the light compared to the *CLASP*_*pro*_:*GFP-CLASP* line (Fig. 3A), there was no such increase in the dark. Interestingly, *CLASP* gene expression in the *35S*_*pro*_:*YFP-CLASP* line was about eightfold higher in the dark compared to the light (Fig. 3G). Taken together with the results using the native *CLASP* promoter, these findings indicate that while CLASP protein levels are reduced in dark-grown roots, *CLASP*-specific transcripts accumulate.

Recognizing that hypocotyl expansion is stimulated in the dark, we measured *CLASP* transcript and protein levels in hypocotyls. In rapidly growing dark-grown hypocotyls, GFP-CLASP remained abundant and could be seen distributed along MTs (Supplemental Movie S1). qRT-PCR from dark-grown seedlings revealed that *CLASP* expression was ∼ 30% higher in the hypocotyls compared to root tips (Supplemental Fig. S2). This demonstrates that in the dark-grown hypocotyls, both GFP-CLASP protein *CLASP* transcript levels remain high, in contrast to the dark-grown root tips, where transcript levels increased 2.5-fold (Fig. 2) but protein levels fall (Fig. 3B). To determine if this uncoupling of transcript and protein levels is a general phenomenon for BR-associated components, we measured the expression of *DWF4*, which encodes a BR biosynthetic enzyme. In contrast to the 30% lower expression of *CLASP* in the root tip compared to the hypocotyl, we found that that *DWF4* expression was reduced by ∼ 80% (Supplemental Fig. S2).

### The CLASP protein is not degraded via the proteasome or autophagy in the dark

To investigate the possibility that CLASP protein was rapidly degraded in the dark, we inhibited two pathways involved in protein degradation and assessed protein abundance using the GFP-CLASP fluorescent reporter (Fig. 4). Plants were treated with 50 µM MG132 for 3 hours and 1 µM concanamycin A for 6 hours to inhibit the 26S proteasome and autophagy, respectively. No increase in GFP-CLASP was observed with MG132 (Fig. 4A, B). As a positive control, we observed PIN2-GFP localization after treatment with 50 µM MG132 for 3 hours because this protein is known to be degraded by the 26S proteasome pathway (Laxmi et al. 2008). PIN2 fluorescence increased at the plasma membrane and decreased in the vacuole, demonstrating that the MG132 drug treatment was functioning as expected (Fig. 4C). GFP-CLASP levels were also unaffected by Concanamycin A treatments (Fig. 4D-E). These results demonstrate that CLASP is not actively degraded in the dark, which suggests instead that the protein levels are regulated by suppression of translation.

**Figure 4.**
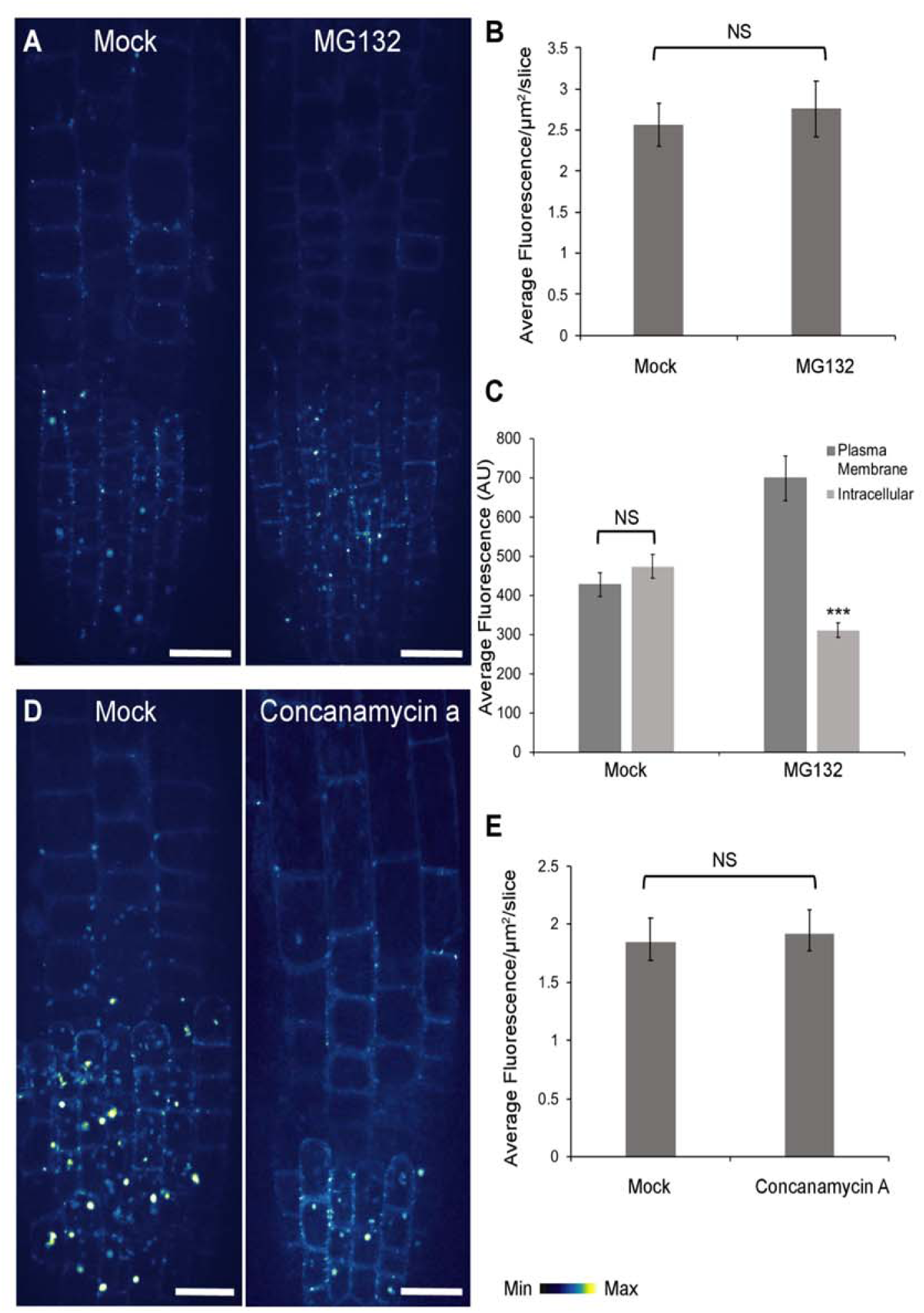
CLASP is not degraded by the proteasome or autophagy in dark-grown roots A, Confocal z-projection of *CLASP*_*pro*_:*GFP-CLASP/clasp-1* plants treated with either a mock solution or 50 μM MG132 for 3h. Scale bar = 10 μm. B, Quantification of mean GFP-CLASP fluorescence intensity in mock and MG132-treated plants shown in (A). NS = no significant difference as assessed by a Student’s t-test with equal variance. n = 20 roots for each treatment. C, Quantification of average PIN2-GFP plasma membrane and intracellular fluorescence intensity under the same treatment conditions in (A) and (B). NS = no significant difference, *** p<0.001 assessed by a Student’s t-test with unequal variance for each treatment. n = 65 cells for mock, n = 58 cell for MG132. D, Confocal z-projection of *CLASP*_*pro*_:*GFP-CLASP/clasp-1* plants treated with either a mock solution or 1 μM Concanamycin A. Scale bar = 10 μm. E, Quantification of mean GFP-CLASP fluorescence intensity in mock and concanamycin A-treated plants shown in (D). NS = no significant difference as assessed by a Student’s t-test with equal variance. n = 20 roots for each treatment.

### Microtubules do not form transfacial bundles in dark-grown meristems

Since CLASP binds to MTs and influences their organization, we hypothesized that CLASP-dependent MT arrays would be disrupted in dark-grown roots. In roots grown in the light, CLASP mediates the formation of transfacial MT bundles that are specific to cells in the division zone (Fig. 5A, B, C). Transfacial MT bundles were completely absent in dark-grown root tips and instead, MTs displayed a predominately transverse array characteristic of cells in the *clasp-1* knockout mutant (Fig. 5D, E, F). In the elongation zone of light-grown roots, MTs are organized transverse to the axis of growth. In contrast, the MTs in dark-grown elongating root cells were oriented along the long axis of the cell (Fig. 5G, Supplemental Movie S2). When increasing amounts of sucrose were added to the growth medium of dark-grown plants, the angle of MT alignment in division-zone cells had a tendency towards more longitudinal than transverse, but was not fully restored to light-grown conditions (Fig. 5H). The addition of sucrose did not have a significant effect on MT alignment in light-grown seedlings (Fig. 5H). These results suggest that MTs have a dramatically different organization when roots are not actively growing in the dark and that sucrose can slightly change MT alignment.

**Figure 5.**
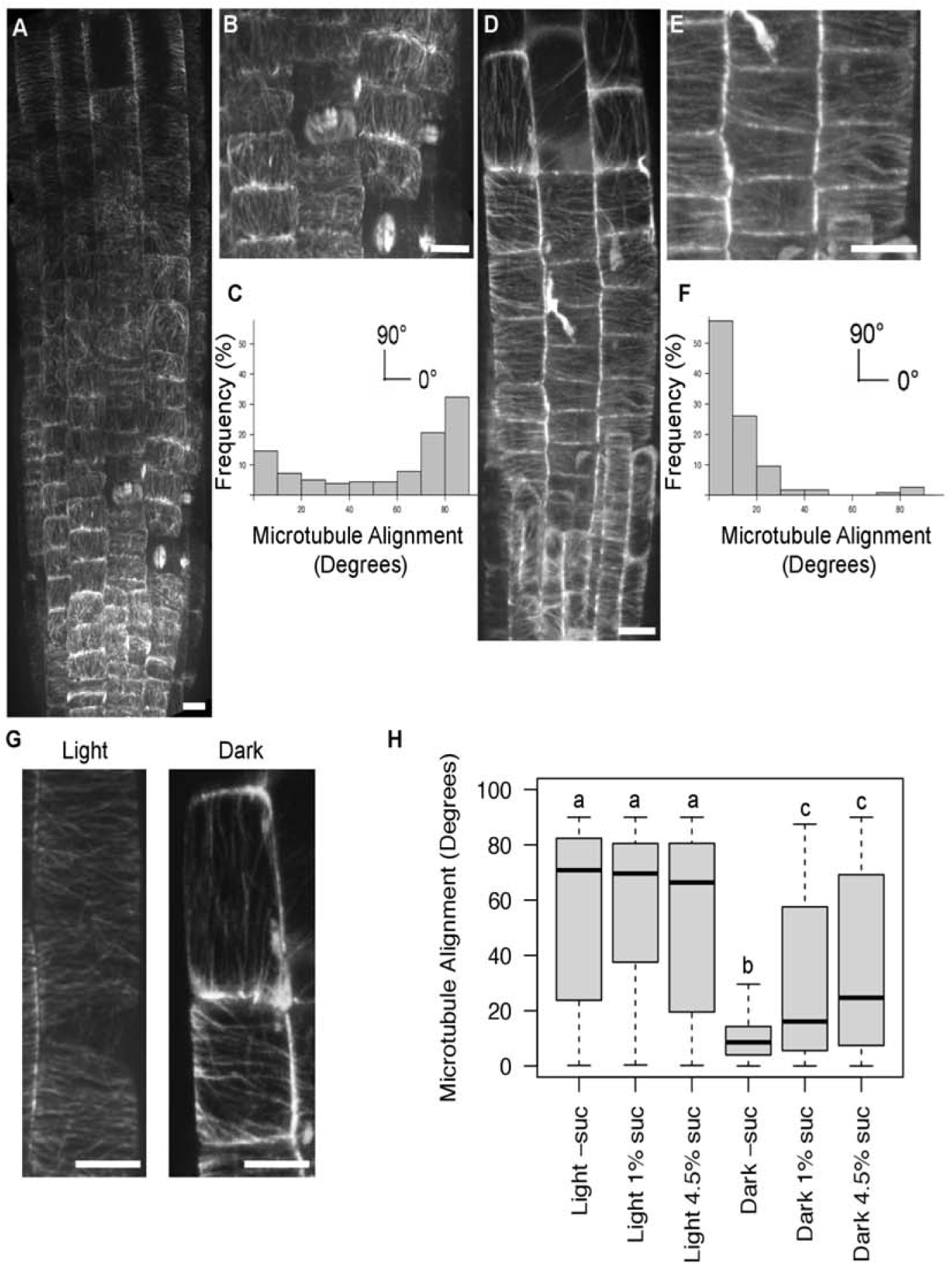
Microtubules (MTs) in the root have different array organizations in the absence of light. A, MTs labelled with GFP-MBD driven by a Ubiquitin promoter in the division zone of the root apical meristem grown in light conditions. Scale bar = 5 μm. B, Higher magnification image of transfacial MT bundles in the division zone of a light-grown root. Scale bar = 5 μm. C, Distribution frequency of MT orientation angles from cells in the division zone of light-grown plants. A MT at zero degrees is transverse; 90 degrees is longitudinal. n = 500 cells from 10 roots. D, MTs in the division zone of the root apical meristem grown in the dark. MTs were labelled with GFP-MBD driven by a Ubiquitin promoter (same fluorescent marker as in A, B). Scale bar = 5 μm. E, Higher magnification image of MTs in cells shown in D. Scale bar = 5 μm. F, Distribution frequency of MT orientation angles from cells in the division zone of dark-grown plants. n = 143 cells from 10 roots. G, GFP-MBD-labelled MTs in elongating cells of plants grown in the light and dark. Scale bar = 5 μm. H, Boxplot of MT orientation in division zone cells of roots grown in either the light or dark on varying amounts of sucrose. The dark horizontal lines mark the median, the box is the range from the 25^th^ to 75^th^ percentile, and the whiskers show the highest and lowest data points within 1.5 times the interquartile range. Different letters indicate significance (p < 0.001) assessed by the Mardia-Watson-Wheeler test. N > 140 cells from at least 10 independent roots for each treatment.

In a previous study, gibberellins (GAs) were identified as regulators of MT organization through the interaction of DELLA proteins and the prefoldin complex, which promotes proper tubulin folding (Locascio et al., 2013). This work showed that when GA is absent, the prefoldin complex is sequestered in the nucleus and decreases the amount of free tubulin heterodimers available for incorporation into the MT. Treatment with GA results in degradation of the DELLA proteins and the movement of the prefoldin complex to the cytoplasm. Based on the sparse MT population observed in dark-grown roots, we hypothesized that GA treatment would induce the prefoldin complex to become functional in the cytoplasm and increase the assembly of MTs, which in turn would result in greater accumulation of CLASP at cell edges. In all cases examined, application of gibberellic acid did not affect MT density or organization (Supplemental Fig. S3). This suggests that the lack of transfacial MT bundles and CLASP protein cannot be rescued by a more active prefoldin complex in dark-grown roots.

### Auxin transporters and BR receptors accumulate in large central vacuoles in dark-grown roots

Previous work has shown that CLASP promotes recycling of the BRI1 receptor (Ruan et al., 2018) and PIN2 transporter (Ambrose et al., 2013) to the plasma membrane. Since CLASP protein levels were depleted in dark-grown seedlings, we hypothesized that the distribution and abundance of BRI1 and PIN2 would be affected in light-versus dark-grown seedlings. We first noticed the formation of large central vacuoles in meristematic cells of dark-grown seedlings, which were absent from cells of the same developmental stage exposed to light (Fig. 6 A-D, brightfield images). GFP-tagged BRI1 and PIN2 were then observed in light and dark conditions (Fig. 6 A-D). In the light treatments, fluorescence intensity was highest at the plasma membrane, both with and without sucrose (Fig. 6A, B, Supplemental Fig. S4). For seedlings grown in the dark with 1% sucrose, PIN2-GFP and BRI1-GFP were still detected at the plasma membrane, although the fluorescence intensity was considerably reduced (Supplemental Fig. S4). In dark conditions in the absence of sucrose, both BRI1 and PIN2 were localized to the vacuoles, indicating that these proteins were undergoing degradation (Fig. 6C, D). Additionally, DR5:GFP labelling in light- and dark-grown roots showed an accumulation of auxin in the meristem of dark-grown roots compared to light-grown roots (Fig. 6E). The transport of BRI1 and PIN2 to the vacuole in dark-grown plants, as well as the pooling of auxin in the root tip, is consistent with reduced recycling of proteins to the plasma membrane when CLASP levels are depleted.

**Figure 6.**
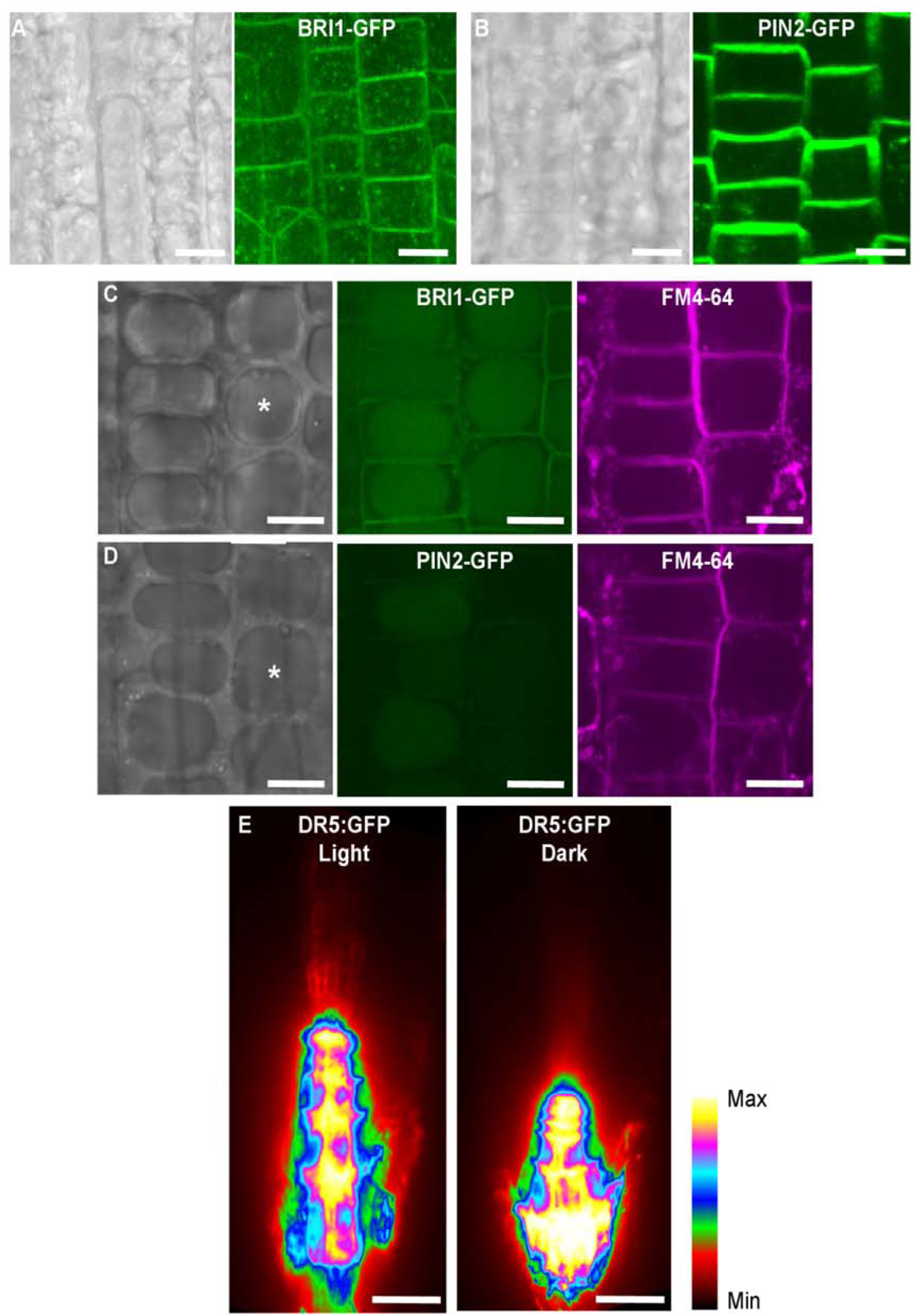
PIN2 and BRI1 are not maintained at the plasma membrane in dark-grown plants. Representative images are confocal z-projections. A and B, BRI1-GFP (A) and PIN2-GFP (B) in light-grown division zone cells of the root. Brightfield images are shown to the right of fluorescence images. Scale bar = 5 μm. C and D, BRI1-GFP (C) and PIN2-GFP (D) in dark-grown division zone cells of the root. Note the large central vacuoles (labelled with asterisks in brightfield images) that are absent in light-grown samples. FM4-64 staining was used to show the plasma membrane and the tonoplast of the vacuole. Scale bar = 5 μm. E, DR5:GFP in light and dark-grown root tips. Scale bar = 30 μm.

## DISCUSSION

Root meristem growth is essential for plant development, and can be readily adjusted based on light availability, nutrients, or abiotic stress. Here, we show how modulation of CLASP protein levels are coincident with either rapid root growth in light conditions, or a cessation of growth in dark conditions (Fig. 7). We showed through fluorescence imaging that BZR1, a negative regulator of *CLASP*, is degraded in dark-grown root meristems. As expected, due to the loss of negative regulation by BZR1, *CLASP* transcript levels were increased in dark-grown root tips. Despite the two-and-one-half fold-elevated transcript level, we detected greatly reduced CLASP protein abundance both when *CLASP* was expressed under its endogenous promoter or overexpressed with the 35S promoter. We also determined that CLASP is not actively degraded in dark-grown roots, leading us to conclude that there is a ‘translational checkpoint’ that maintains low levels of CLASP when root cell division and elongation are not necessary, such as in the dark when plants must prioritize hypocotyl expansion. It is important to note that in early seedling development and prior to becoming photosynthetically active, plants must ration sugar reserves. Our work also clearly demonstrates that supplementing growth media with sucrose while growing plants in the dark obscures the dramatic phenotypes associated with cellular starvation, which plants would likely experience when photosynthetic output is limited.

**Figure 7.**
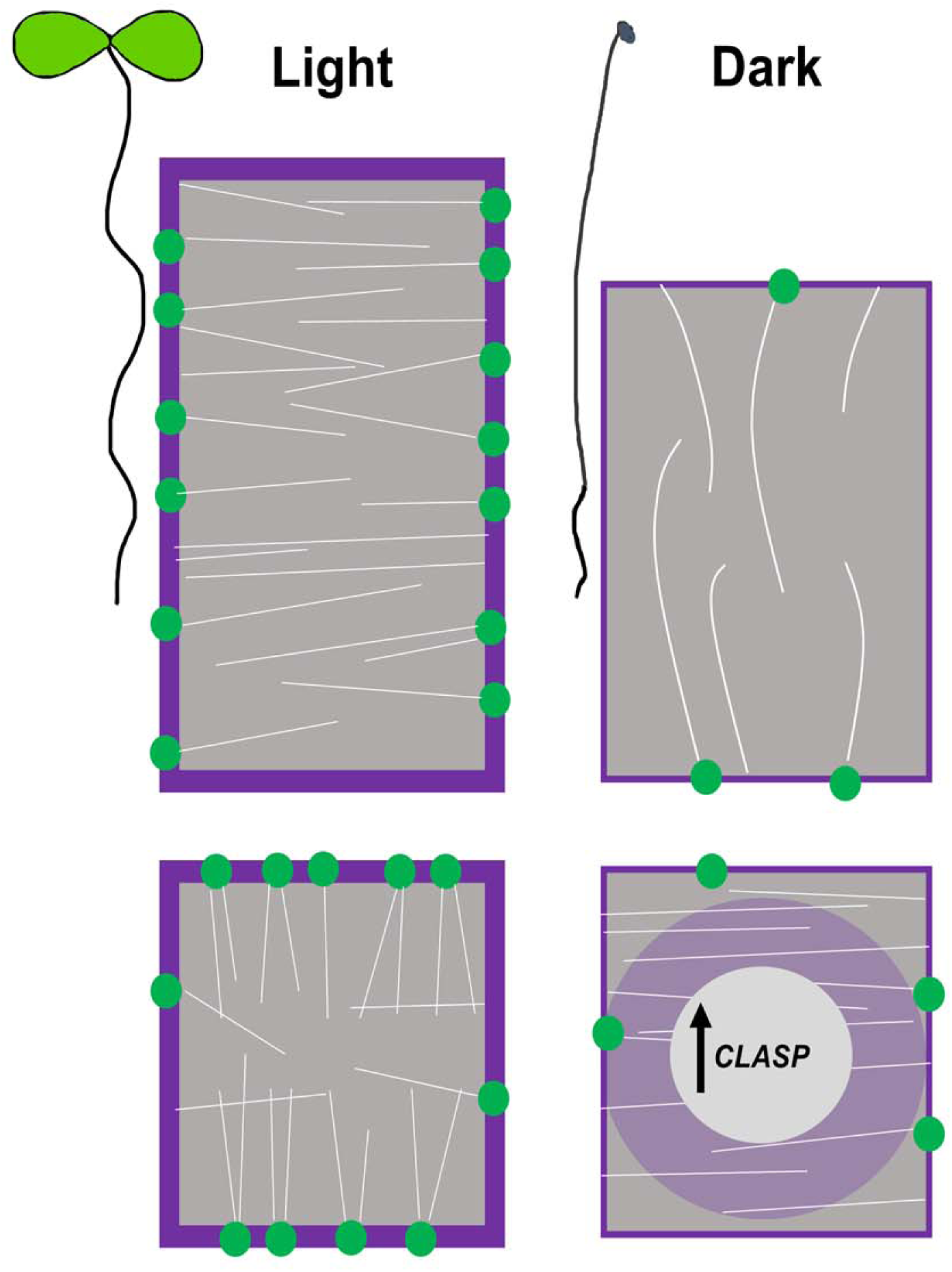
Summary schematic to illustrate how CLASP is implicated in the response to light or dark growth conditions. CLASP (green dots) localizes to the sharp transverse cell edges in dividing cells and mediates the formation of transfacial MT bundles (white lines). CLASP also fosters the recycling of PIN2 and BRI1(purple) to the plasma membrane. In elongating cells, CLASP is found along the longitudinal edges and MTs display a transverse pattern. In sucrose-deprived roots grown in the dark, CLASP levels are diminished, resulting in the disappearance of MT bundles and PIN2 and BRI1 in the large central vacuole. *CLASP* transcript levels remain high in the dark, suggesting a translational mode of regulation.

Our study has identified a very specific and specialized regulatory system that controls CLASP protein levels in response to light/dark conditions, and further underlines the importance of CLASP as a regulator of meristem size in the *Arabidopsis* root tip (Ambrose et al., 2007). *CLASP* was previously shown to be transcriptionally downregulated by the BR signalling pathway (Ruan et al., 2018), but regulation at the translational level has not yet been demonstrated. Our results indicate a general dampening of the BR signalling pathway in root meristems under dark conditions, which is consistent with a study showing BZR1 degradation when carbon availability was limited (Zhang et al., 2016), and another study that found *DWF4* transcript was reduced in dark-grown root tips (Sakaguchi and Watanabe, 2017). Like *CLASP, DWF4* is also suppressed by BZR1 (He et al., 2005) but does not show the same effect of increased transcript as *CLASP* in dark-grown root meristems. Transcript accumulation under these conditions therefore does not appear to be a ubiquitous response, since genes such as *BRI1, DWF4, MOR1*, and the mitosis checkpoint regulator *CYCB1* showed low expression compared to light-grown root tips. In the dark, GFP driven by the *CLASP* promoter showed high levels of fluorescence intensity compared to the light, suggesting that free GFP, which does not have a function in plants, is readily translated whereas the CLASP protein is not. Live cell imaging confirmed that GFP-CLASP was abundant along MTs in expanding etiolated hypocotyls, suggesting that translational regulation of CLASP is organ-specific.

When CLASP protein levels were diminished in dark-grown root tips, MT organization was impacted. Transfacial MT bundles did not form and cells displayed transverse MT arrays, similar to the *clasp-1* knockout mutant. The role of transfacial MT bundles in root interphase cells is currently unknown, but these arrays are specific to proliferating cells and dependent on CLASP. The MTs in dark-grown cells were sparse, suggesting that the overall amount of tubulin was reduced. MTs in elongated cells of dark-grown roots displayed a longitudinal pattern, which is in stark contrast to the hyperparallel transverse array seen in rapidly elongating cells. SCAR1, a component of the actin-nucleation complex, is degraded by the COP1 E3 ligase and 26S proteasome in the dark, leading to the loss of longitudinal F-actin and reduced root growth (Dyachok et al., 2011). It remains to be determined if components of the root MT cytoskeletal machinery are affected in a similar manner by growing plants in the dark.

When plants were deprived of light and a source of sucrose, the auxin transporter PIN2 and BR receptor BRI1 were less abundant at the plasma membrane and accumulated in the vacuole. This redistribution is consistent with a lack of CLASP, which normally promotes recycling of PIN2 and BRI1 to the plasma membrane through tethering of SNX1 vesicles (Ambrose et al., 2013; Ruan et al., 2018). Cells in the division zone of roots normally have several small vacuoles, whereas an enlarged central vacuole is characteristic of elongating and differentiated cells. It may be that cell wall properties play a role in vacuolar expansion in these cells, as mutants of extracellular leucine-rich repeat extensins and the receptor-like kinase FERONIA displayed enlarged vacuoles in root meristematic cells (Dünser et al., 2019), similar to what we observed in our study.

What mechanism controls translation in *Arabidopsis* root tips during carbon-limited conditions? Plants maintain elaborate nutrient-sensing signalling pathways that impinge on transcriptional and translational network remodelling. Carbon derived from the shoot activates TOR signalling to induce meristem growth in the root. In addition to activating genes related to amino acid synthesis, the cell cycle, and RNA synthesis and processing, TOR phosphorylates the E2Fa transcription factor that activates S-phase related genes (Xiong et al., 2013). When these processes are inhibited under nutrient starvation, mature ribosomes are degraded by autophagy (a process termed ribophagy) that has so far been described in yeast (Kraft et al., 2008) and mammals (An and Harper, 2018; Wyant et al., 2018). Work on ribophagy has been limited in plants, although Floyd et al. (2015) demonstrated rRNA degradation in the vacuole in a ribonuclease mutant even under sufficient nutrient conditions. If ribophagy is occurring in dark-grown root meristems, it is possible that the buildup of *CLASP* mRNA is due to a reduction in translational machinery to efficiently process transcripts.

Plants must ensure a strategic partitioning of resources to survive before they can become photosynthetic and produce sugar. The minimal levels of CLASP protein and other components necessary for cell proliferation are consistent with an inhibition of division in the root meristem when nutrients are limited. It remains unknown if this CLASP-specific response occurs in other systems. The only other evidence is from fission yeast, where upon glucose starvation, the CLASP homologue Cls1 was negatively regulated by Protein Kinase A, causing MT destabilization and delayed cell division (Kelkar and Martin, 2015). Since GFP-CLASP fluorescence did not increase when protein degradation mechanisms were inhibited, it is not likely that CLASP is being synthesized and degraded, which would be a costly energetic investment. It may be advantageous for plants to accumulate *CLASP* transcripts so that root growth can be readily promoted under the appropriate environmental circumstances. Given that plants undergo daily cycles of light and dark exposure, it would be of interest to determine if these transcriptional, translational, and cellular modifications follow a diurnal pattern. The regulation of *CLASP* in response to environmental signals supports a role for this gene in modulating root growth when plants are subjected to stress during their development.

## MATERIALS AND METHODS

### Plant Material and Growth Conditions

*Arabidopsis thaliana* plants used in this study were all from the Columbia-0 ecotype background. Seeds were surface sterilized in a solution of 50% v/v ethanol and 3% v/v hydrogen peroxide and grown on solid ½ MS medium (pH 5.8) with 1% bacto agar and different concentrations of sucrose as indicated in the experimental data. The seeds were kept in the dark at 4°C for 2-3 days, then germinated in growth chambers under continuous light at 22°C for 6 days. For dark-grown seedlings, the samples were exposed to 8 h of light to promote germination, after which the petri dishes were wrapped in foil for the remainder of the growth period.

The *CLASP*_*pro*_:*GFP-CLASP* construct fully rescued the *clasp-1* null mutant as described in (Ambrose et al., 2011). The Cytrap marker lines were kindly given to us by Dr. Masaaki Umeda and are described in (Yin et al., 2014). The BZR1-CFP lines were given to us by Dr. Zhi-Yong Wang and are described in (Wang et al., 2002). The *35S*_*pro*_:*YFP-CLASP* reporter was described in (Kirik et al., 2007) and given to us by Dr. Viktor Kirik. Microtubules were visualized with lines expressing *UBQ1*_*pro*_:*GFP-MBD* (Ruan et al., 2018). The *BRI1*_*pro*_:*BRI1-GFP* was from Dr. Joanne Chory and described in (Geldner et al., 2007). *PIN2*_*pro*_:*PIN2-GFP* is described in Xu and Scheres, (2005) and *DR5:GFP* in Benková et al. (2003).

### Propidium Iodide Staining

To quantify meristem cell number, root tips were stained in a 10 μg/mL solution of propidium iodide dissolved in distilled water for 1 minute. This was followed by three 1 minute rinses in dH_2_O before mounting on a glass slide for microscopy.

### Confocal Microscopy

6-day-old seedlings were mounted in either 1/2 MS (no sucrose) liquid or drug solutions on glass slides for imaging. Fluorescent protein reporters were imaged using a spinning-disk confocal microscope (Leica DMi8 inverted microscope, Perkin-Elmer UltraView spinning-disk system, and a Hamamatsu 9100-02 Camera) with either a 40x/NA 1.25 oil or 63x/NA 1.3 glycerol lens. CFP was imaged with a 440 nm laser and 480/40 emission filter, GFP was detected using a 488 nm laser and 525/36 emission filter, and YFP was imaged using a 514 nm laser and 540/30 emission filter. For propidium iodide and FM4-64, excitation was with a 561 nm laser and 595/50 emission filter. The slice thickness for z-projections was 0.3 μm. Images were captured using the Volocity 6.3 software package (Perkin Elmer).

### Image Analysis

Confocal images were processed using ImageJ (https://imagej.nih.gov/ij/). Root length was quantified using the NeuronJ plugin (Meijering et al., 2004). Fluorescence intensity of GFP-CLASP in root meristems was measured by drawing regions that encompassed epidermal cells from the quiescent centre to the beginning of the elongation zone from compressed z-stacks. The “corrected fluorescence” was calculated by subtracting the mean background fluorescence and then dividing by area of the measured region and number of z slices to facilitate comparisons between light and dark samples. Microtubule orientation within meristematic cells was calculated using the ImageJ plugin FibrilTool (Boudaoud et al., 2014). BRI1 and PIN2 fluorescence was measured in two cell files from six independent roots in cells extending from the quiescent centre to the elongation zone. Briefly, the sum slices tool was used to generate z-projections that extended from the top-most part of the cell to 8-9 μm deep. The plasma membrane fluorescence was measured by tracing the area using the segmented line tool (pixel width 3), while the intracellular space fluorescence was measured with the polygon tool.

### RNA Extraction and Quantitative-Real-Time PCR

6-day-old seedlings were used for gene expression analysis. About 400 root tips for each treatment were excised directly beneath the differentiation zone where root hairs were visible (a length of approximately 2 mm and 0.5 mm for light and dark-grown roots, respectively. The root tips were ground in TRIZOL reagent (Invitrogen, Life Technologies) and RNA was extracted. The quality of the RNA was assessed on an agarose gel before proceeding. Samples were treated with DNaseI (Amplification Grade, Invitrogen) to remove any residual DNA, and then subjected to reverse transcription with SuperScript III Reverse Transcriptase (Invitrogen, Life Technologies) to obtain cDNA. qRT-PCR was carried out with SensiFAST(tm) SYBR® & Fluorescein Mix (BIOLINE) in a Bio-Rad iQ5 thermal cycler. The Pfaffl method (Pfaffl 2001) was used to calculate relative expression levels. The primers used are listed in Supplemental Table S1.

### Gibberellic Acid Treatment

Plants were grown in the dark for 6 days as described in the Plant Material and Growth Conditions section. The samples were either germinated on media containing 10 μM GA for 6 days, or grown on ½ MS (no sucrose) media for 5 days, then transferred to media containing 10 μM GA for 24 h. The mock treatment was 0.1% ethanol.

### Drug Treatments

Seedlings were transferred to ½ MS medium containing 50 μM MG132 (Sigma-Aldrich) for 3h or 1 μM Concanamycin A (Sigma-Aldrich) for 6h to inhibit the proteasome and autophagy, respectively. The mock treatment was ½ MS medium with 0.5% DMSO.

### Statistical Analyses

Statistical analyses were performed in R (v. 3.3.3; The R Foundation for Statistical Computing, 2017). The Mardia-Watson-Wheeler test was performed using the Circular Statistics package in R (https://r-forge.r-project.org/projects/circular/).

## Supplemental Data

The following supplemental materials are available.

**Supplemental Figure S1**.

Confocal z-projection of wild-type dark-grown root meristems showing autofluorescence with GFP laser settings (left image) and brightfield channel (right image). Scale bar = 30 μm.

**Supplemental Figure S2**.

*CLASP* and *DWF4* transcript levels in the hypocotyl and root of dark-grown plants. *APT1* was used as the reference gene. Error bars denote SE. n = 3 biological replicates.

**Supplemental Figure S3**.

Gibberellic acid treatment does not change microtubule organization in dark-grown root meristems. GFP-tubulin labelled microtubules in root meristems treated with mock (0.1% ethanol) or GA for either 24 h or 6 days. Sale bar = 20 μm.

**Supplemental Figure S4**.

BRI1 and PIN2 localization in root epidermal cells of light and dark-grown plants with 1% sucrose. A and B, BRI1-GFP grown in the light (A) and dark (B) with 1% sucrose. C and D, PIN2-GFP grown in the light (C) and dark (D) with 1% sucrose. Scale bars = 30 μm

**Supplemental Movie S1**

Time-lapse of GFP-CLASP in a 6-day-old etiolated hypocotyl. Related to Fig. 3.

**Supplemental Movie S2**

Time-lapse of microtubules labelled with GFP-tubulin in elongating cells of dark-grown roots. Related to Fig. 5.

**Supplemental Table S1**

Primer sequences used for qRT-PCR analysis.

## ACKNOWLEDGMENTS

This work was supported by an NSERC Discovery grant (RGPIN-2019-05432), the Canada Research Chairs Program, and the Canada Foundation for Innovation to G.O.W, as well as an NSERC CGS-D grant to L.H. Microscopy was conducted in the UBC Bioimaging Facility.

